# New insights on the role of HCN in root hair elongation through single cell proteomics

**DOI:** 10.1101/2022.01.05.475094

**Authors:** Lucía Arenas-Alfonseca, Masashi Yamada, Luis C. Romero, Irene García

## Abstract

Root hairs are specialized structures involved in water and nutrient uptake by plants. They elongate from epidermal cells following a complex developmental program. ß-cyanoalanine synthase (CAS), which is mainly involved in hydrogen cyanide (HCN) detoxification in *Arabidopsis thaliana*, plays a role in root hair elongation, as evidenced by the fact that *cas-c1* mutants show a severe defect in root hair shape. In addition to root hairs, CAS C1 is expressed in the quiescent center and meristem. However, the *cas-c1* mutation has no visible effect on either tissue, in both control and nutrient-deprivation conditions. To identify its role in root hair formation, we conducted single cell proteomics analysis by isolating root hair cells using Fluorescence-Activated Cell Sorting (FACS) from wild type and *cas-c1* mutants. We also analyzed the presence of *S*-cyanylation, a protein post-translational modification (PTM) mediated by HCN and affecting cysteine residues and protein activity, in proteins of wild type and *cas-c1* mutants. We found that several proteins involved in root hair development, related to the receptor kinase FERONIA signaling and to DNA methylation, are modified by this new post-translational modification.

**One sentence summary:** Arabidopsis root hair proteomics unveals that several proteins involved in root hair development are susceptible of modification by *S*-cyanylation.

## Introduction

Root formation is one of the most important evolutionary events in the adaptation of plants to terrestrial ecosystems. Root hairs are specialized tubular structures of epidermal cells that improve the incorporation of nutrients and anchoring of the plant to the soil, as well as playing an important role in the interaction with other organisms (Cui et al., 2018).

In Arabidopsis, epidermal cells present a specific distribution that determines their fate: those that are between two cortical cells will become root hairs (H cells), and those in contact with the membrane of a single cortical cell will develop into non-hair (N) cells (Datta et al., 2011). The differentiation of N or H cells is determined by a plethora of regulatory proteins, which have been extensively studied (Grierson et al., 2014). Once cell fate is determined, H cells change their shape by modifying the cell wall to allow for root hair elongation. To accomplish this, ROP (“RHO-RELATED GTPase”) proteins are recruited by the FERONIA (FER) receptor kinase (Duan et al., 2010) and are concentrated in the root hair growth zone (Molendijk et al., 2001). At the same time, endoplasmic reticulum and filamentous actin are concentrated in this area. In addition, the pH decreases to 4-4.5, activating expansins and allowing the relaxation of the cell wall and the formation of the initial bulge (Monshausen et al., 2007). Elongation of the root hair is directed by vesicles produced by the rough endoplasmic reticulum and the Golgi complex, which release polysaccharides and cell wall glycoproteins, as well as protein synthases and membrane transporters through exocytosis at the tip of the root hair (Ketelaar et al., 2008). The control of membrane trafficking, therefore, is essential in the development of root hairs. RAB GTPases such as RAB-A4b (Rab GTPase Homolog A4b) play an important role in the regulation of vesicular traffic (Nielsen et al., 2008) by controlling the phosphorylation state of phosphatidyl inositol (PI) -PI4P (Thole et al., 2008) (Bubb et al., 1998; Yoo et al., 2012). Another ROP GTPase, ROP2 (Rho-related protein from plants 2) initiates ROS production by NADPH oxidase RHD2/ATRBOHD (Root hair defective 2) (Foreman et al., 2003; Jones et al., 2007). ROS promotes the entry of Ca^2+^, which activates RHD2 in a positive feedback loop (Mendrinna and Persson, 2015). ROS production is also induced by the RSL4 protein (Root hair defective 6 like 4), which controls the expression of NADPH oxidases and peroxidases. These also participate in the production of ROS in root hairs (Hwang et al., 2017). The Ca^2+^ gradient at the tip of the root hair directs the growth, facilitates the fusion of the vesicles with the apical plasma membrane and contributes to the release of the vesicular content necessary for the expansion of the cell wall (Pei et al., 2012). The organization of the cytoskeleton, hormones such as ethylene and auxin as well as nutrient availability are also essential in the formation and development of root hairs (Ketelaar, 2013) (Sieberer et al., 2005). (Jones et al., 2009) (Cui et al., 2018) (Jungk, 2001).

Hydrogen cyanide (HCN) is synthesized in plants as a co-product during ethylene biosynthesis but also is produced by several plant growth-promoting rhizobacteria. It acts as a biocontrol agent and influences nutrient availability (Peiser et al., 1984; Rijavec and Lapanje, 2016). The role of HCN as a signaling gasotransmitter involved in root hair development and the plant immune response has been widely described. Recently, it has also been described as a signaling molecule in mammalian systems (Garcia et al., 2010; Garcia et al., 2013; Gotor et al., 2019; Pacher, 2021; Zuhra and Szabo, 2021). HCN is mainly detoxified by the ß-cyanoalanine synthase CAS-C1. Maintaining low levels of HCN by CAS-C1 is required for proper root hair development (Garcia et al., 2010). CAS-C1 accumulates in root hair tips during root hair elongation. Thus, higher accumulation of HCN in *cas-c1* inhibits the action of the NADPH oxidase gene *ROOT HAIR DEFECTIVE 2 (RHD2)/AtrbohC* in a manner independent of enzyme inactivation (Arenas-Alfonseca et al., 2018b, 2018a). Root hairs of *cas-c1* are specified correctly, but they do not elongate. Genetic analysis showed that the mutant of the *SUPERCENTIPEDE* (*SCN1*) gene is epistatic to *cas-c1* and *rhd2* (Arenas-Alfonseca et al., 2018b, 2018a). HCN acts as a signaling molecule that influences root hair development in a manner independent of ROS formation and of direct NADPH oxidase inhibition, and its action has been localized in the first steps of root hair elongation, between SCN1 action and RHD2-driven production of O_2_·^-^ (Arenas-Alfonseca et al., 2018b, 2018a).

The mechanisms that underlie HCN modulation, the mode of action and the specific targets of this molecule are areas of active research. HCN is able to produce post-translational modifications in the thiol groups of cysteines, producing *S*-cyanocysteine. Our research has shown that some *Arabidopsis* proteins are naturally *S*-cyanylated and that this modification can change the activity of these proteins (Garcia et al., 2019) suggesting that the *S*-cyanylation represents a new mechanism of regulation of the processes in which HCN plays a signaling role.

Although CAS-C1 is expressed in the meristem in addition to root hairs, we show that *cas-c1* does not affect meristem formation or root hair cell fate. In order to identify the protein targets of cyanide in the root hair, we carried out a cell-type specific proteomic approach to characterize the proteins present in wild-type root hair cells and compare them to those present in *cas-c1* mutant cells. Through this analysis, we found that several proteins involved in root hair development are modified by *S*-cyanylation.

## Results

### *CAS-C1* transcription is cell-type specific

A cell-type specific transcriptomic analysis (Li et al., 2016b), showed that *CAS-C1* is highly expressed in several root cell types, such as the quiescent center (QC, *WOX5*-tagged cells), the phloem pole pericycle (S17-tagged cells), developing cortex (*CO_2_*-tagged cells) and mature cortex (*COR*-tagged cells) (Supplemental Fig. S1A). Also, *CAS-C1* transcription increased in the elongation zone compared to the meristematic zone, and remained high in the differentiation zone (Supplemental Fig. S1B). These results suggested a possible developmental role of CAS-C1 in the root of *Arabidopsis thaliana*.

### *CAS-C1* is not involved in QC formation, meristem length or root elongation

To determine if *cas-c1* mutants are affected in QC establishment or maintenance, we carried out confocal microscopy of wild type and *cas-c1* mutant meristems stained with propidium iodide to visualize cell walls. Fig. 1 shows that no differences were found either in the number or morphology of QC cells in roots grown in control conditions. No differences were also observed in meristem size of wild type and *cas-c1* (white arrows in Fig. 2A mark the transition zone between meristematic and elongation zones). Time-lapse measurements of root length (Fig. 2 and Supplemental Fig. S2) confirmed that root elongation was identical in both genotypes.

**Fig. 1.**
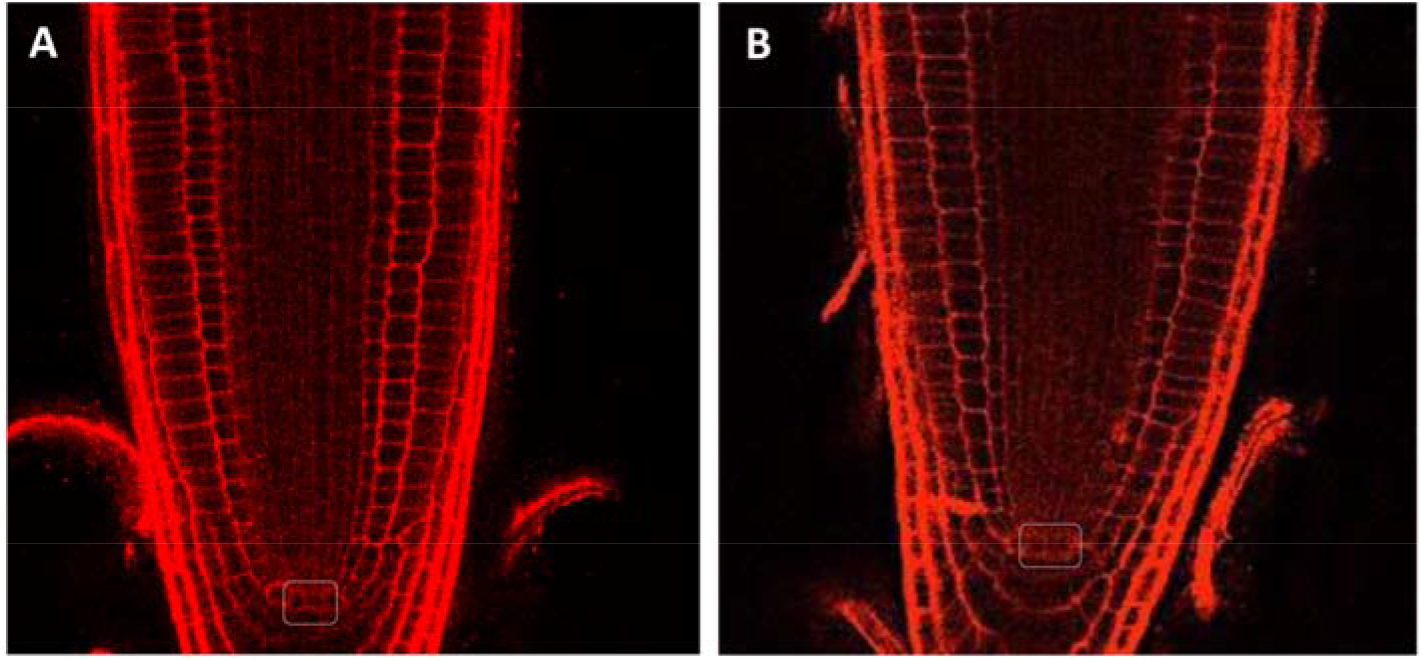
Root meristem phenotype in wild type and *cas-c1* mutant. Col-0 (A) and *cas-c1* mutant (B) seedlings were grown for 7 d on MS medium supplemented with sucrose in vertical plates. Representative images are shown. White squares indicate QC cells. Propidium iodide was used to visualize cell walls.

**Fig. 2.**
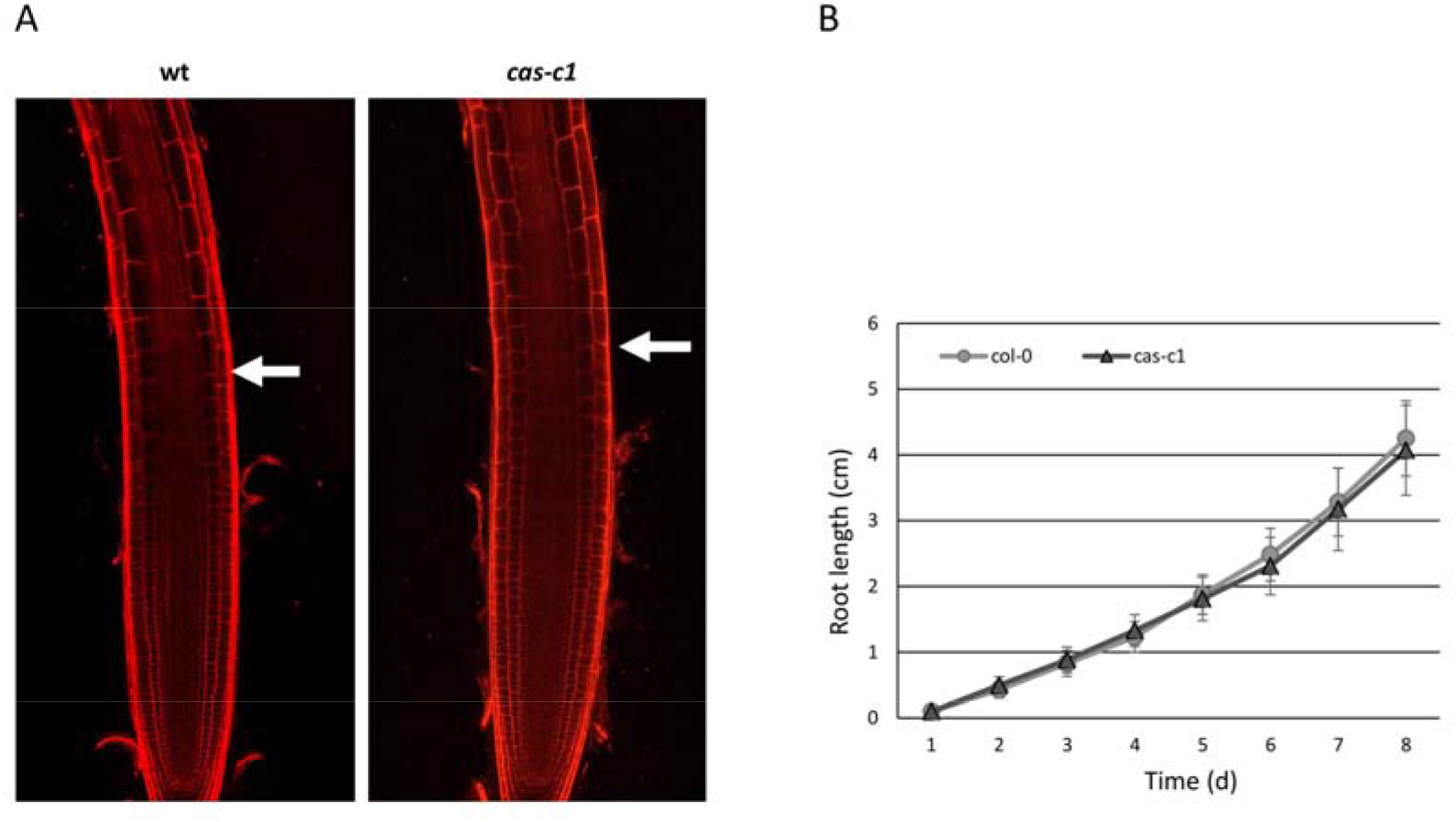
Root meristem size in MS medium. A) A representative root of 7-days-old wild type (wt) and *cas-c1* mutant stained with propidium iodide. White arrows indicate the transition between the elongation and transition zones in wild type and cas-c1 mutant seedlings. B) Root meristem length of wild type and *cas-c1* mutants in MS medium. Media ± SD are shown in the graph. N>20.

The CAS-C1 protein is involved in mitochondria sulfur metabolism. It detoxifies hydrogen cyanide by using as a substrate the sulfur-containing amino acid cysteine (Romero et al., 2014). It is also indirectly involved in nitrogen recycling because the nitrogen residue in HCN can be incorporated into amino acids through the nitrilase activities of ß-cyanoalanine, the product of HCN detoxification by CAS-C1 (Piotrowski, 2008). To determine if root growth is affected by loss of these metabolic functions, the root phenotype was analyzed in conditions of nitrogen and sulfur deficiency. Wild type and *cas-c1* mutant plants were grown on vertical plates of MS medium and different nutrient-deprived media for 9 days and root length was measured (Fig. 3 and Supplemental Fig. S3). No differences were observed in any condition between wild type and *cas-c1* (Supplemental Fig. S4). Phosphate starvation induces, among other root phenotypes, root hair elongation, involving auxin, ethylene, cytokinin and post-translational modifications (Huang and Zhang, 2020). Therefore, as *cas-c1* mutants show a defect in root hair formation, we included phosphate starvation in our observations. Again, no differences were found in Pi starvation (Fig. 3 and Supplemental Figs. S3C and S4D). Taken together our results indicate that *cas-c1* does not affect root growth in control or under N, S or P nutritional stress conditions.

**Fig 3.**
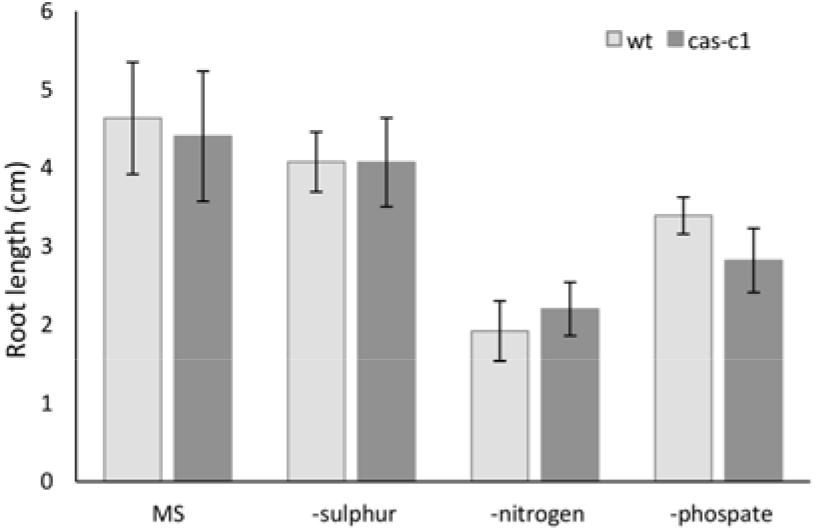
Root length measurement in nutrient deficient media. Wild type and *cas-c1* mutant seedling were grown for 9 d in vertical plates of MS medium or MS medium in the absence of different nutrient compounds. Root length was measured with ImageJ software. Values are means ± SD of three independent experiments.

### *CAS-C1* is not involved in root hair cell fate

Because *cas-c1* has a severe defect in root hair morphogenesis (Garcia et al., 2010; Arenas-Alfonseca et al., 2018a, 2018b), we hypothesized that it may also have altered root hair cell fate. To visualize root hairs during development, we crossed into *cas-c1* the COBRA-like protein 9 (COBL9) promoter driving GFP, which is root hair specific (Brady et al., 2007b) (Petricka et al., 2012). The *cas-c1*; pCOBL9:GFP line showed a normal patterning of green fluorescence (Fig. 4), both in its abnormal root hairs and in its root hair primordia, indicating that the patterning of root hair formation is not altered in this mutant. Thus, *cas-c1* affects root hairs exclusively in their elongation.

**Fig. 4.**
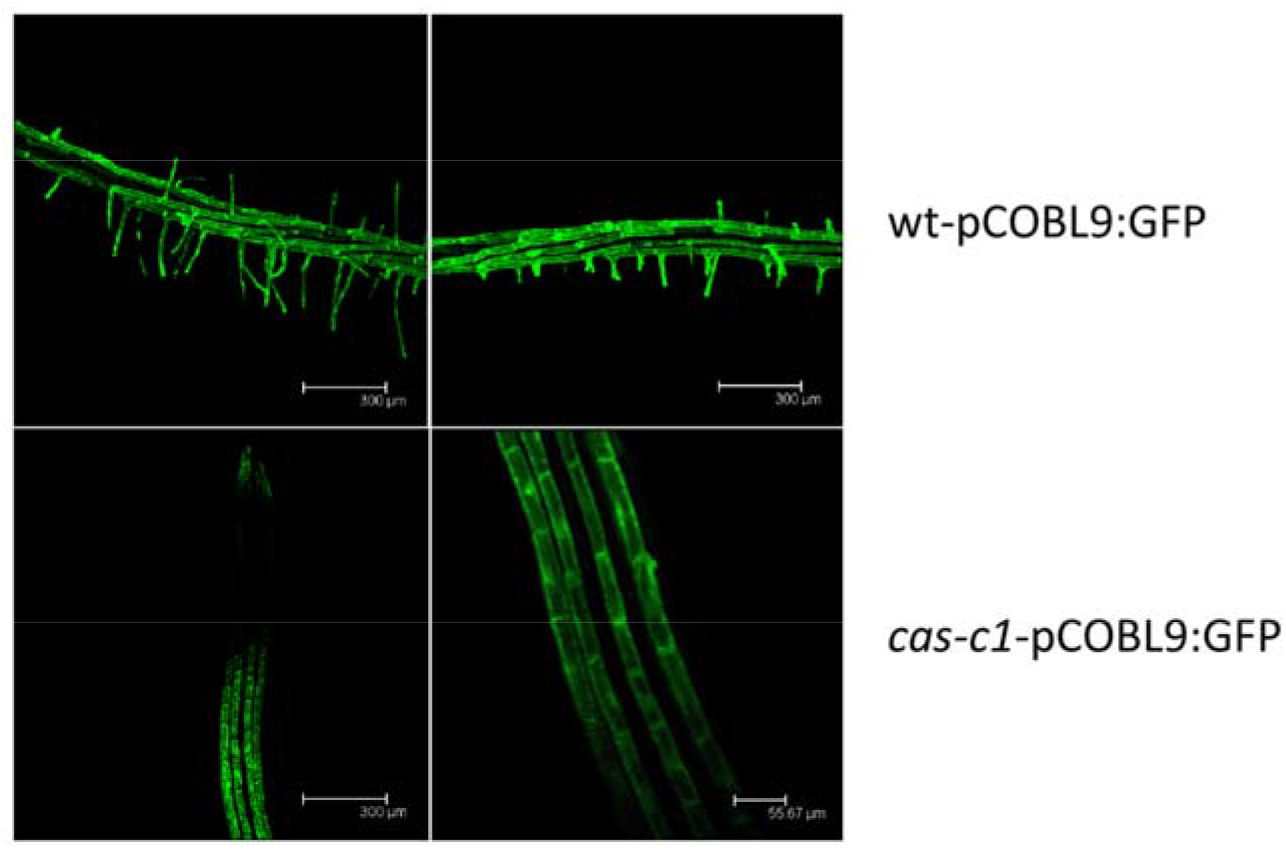
GFP localization in roots from the wt-pCOBL9:GFP and *cas-c1*-pCOBL9:GFP transgenic lines. Plants were grown 6 days in MS medium. Representative images are shown.

### Proteomics of wild-type and *cas-c1* root hairs

To analyze the effect of the *cas-c1* mutation, we isolated 10^6^ root hair cells by Fluorescence-activated cell sorting (FACS) from three independent replicates of WT pCOBL9:GFP and *cas-c1* pCOBL9:GFP roots. The extracted proteins from each sample were trypsin-digested and the peptides analyzed by liquid chromatography-high resolution mass spectrometry (LC-MS/MS).

In wild-type samples, we identified 3,829 unique proteins at a false discovery rate below 1% (FDR<1%), which represent almost 10% of the total *Arabidopsis* proteome (Dataset S1). Previous work identified 1,387 proteins in root hair cells by a similar experimental approach using GeLC-MS/MS analysis (Petricka et al., 2012) (Fig. 5A), 90% of which we also identified. In protein extracts of the *cas-c1* pCOBL9:GFP roots, we identified 3,972 proteins (Dataset S2) of which 3,515 were common in both cell types, 310 were only identified in wild type and 457 only in *cas-c1* (Fig. 5B).

**Fig. 5.**
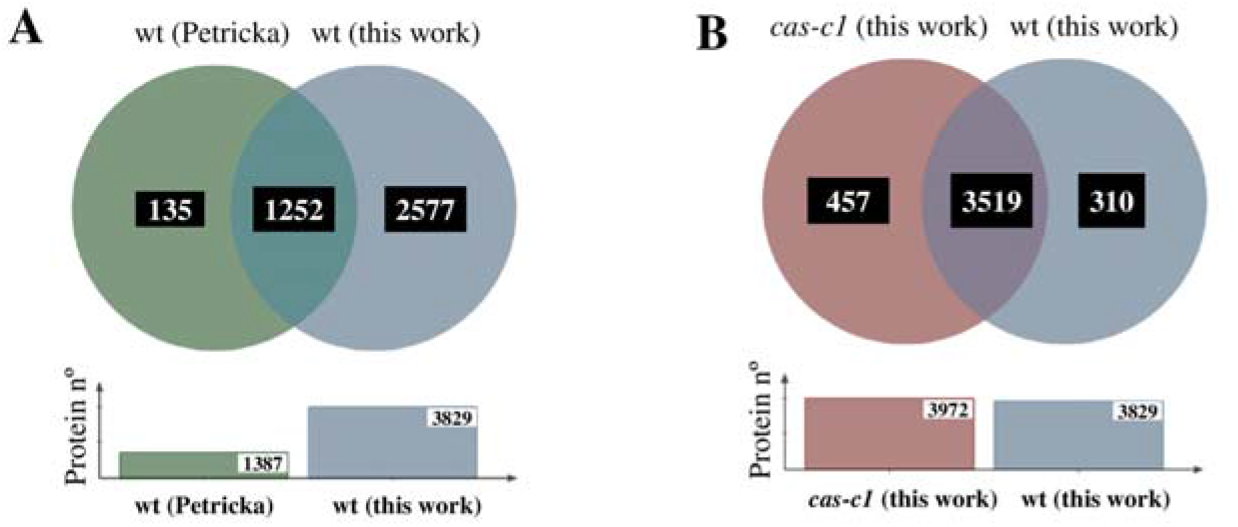
A, Venn diagram showing the unique and intersection of the identified protein in the proteomic analysis of Petricka et al, 2012 and this work. B, Venn diagram showing the unique and intersection of the identified proteins in wild type and *cas-c1* root hair samples.

### *cas-c1* root hair cells differ from non-hair cells at the proteomic level

To understand the stage at which *cas-c1* root hair development is blocked, we compared proteins present in *cas-c1* and in wild-type root hairs. To this end, we compared *cas-c1* and wild-type RH cells with published proteomics data of hair/non hair (NH) epidermal cells (Petricka et al., 2012). Of the 457 proteins present only in *cas-c1* RH and the 113 specific to NH cells, only 5 proteins were coincident (Supplemental Fig. S5). Thus, at the protein level, *cas-c1* RH are not equivalent to NH cells. This is consistent with the *cas-c1* phenotype, which has root hairs, although they are short and unstructured. A similar result was obtained by comparing the proteome of *cas-c1* RH with the wild-type NH proteome in (Lan et al., 2013), in which another RH-specific promoter, EXP7, was fused to GFP and root hair cells isolated by FACS. Thus, these proteomic comparisons show that *cas-c1* root hairs are not similar to non-hair cells and they have their own identity. A deeper analysis of the root hair proteome was therefore undertaken.

### *feronia* and *cas-c1* mutants share common features

Several *Arabidopsis* mutants have a phenotype similar to *cas-c1* including the basic helix-loop-helix (bHLH) transcription factor ROOT HAIR DEFECTIVE 6-LIKE 4 (*rsl4*), which is a target of the bHLH transcription factor ROOT HAIR DEFECTIVE 6 (*rhd6*) and acts as signal to induce cell growth in root hairs (Menand et al., 2007; Yi et al., 2010). We thus compared the transcriptomic data from the genes mis-regulated in *rsl4* (Yi et al., 2010) with the proteins present only in wild-type or in *cas-c1* root hairs. The comparison showed that the number of common genes/proteins was very low and therefore we cannot establish a relationship between theseh mutations (Suppplementary Fig. S6).

RHO GTPases are a large family of related monomeric GTP-binding proteins that serve diverse signaling functions (Etienne-Manneville and Hall, 2002). Plant ROPs are a unique clade of RHO-like GTPases involved in cell growth and polarity establishment among other functions (Yang and Fu, 2007; Kost, 2008). FERONIA is a ROP-interacting receptor-like kinase (RLK) and regulates ROP-signaling and ROS-mediated root hair development, as well as many other aspects of plant growth and development (Hématy and Höfte, 2008). We found that *feronia* (*fer*) mutants show a high percentage of up or down-regulated genes common to the proteins that are present in *cas-c1* root hairs (Supplemental Fig. S7), as well as a very similar phenotype to *cas-c1* at the root hair level (Duan et al., 2010). This suggested a functional relationship between *fer* and *cas-c1* mutations.

### Functional classifications of root hair-specific proteins

To identify the function of CAS-C1 and explain the hairless phenotype of the mutant we addressed the functional classification of the proteins identified in wild-type and *cas-c1* root hair samples based on the information available in three gene/protein databases: Uniprot for Gene Ontology classification (Bairoch et al., 2005; Pundir et al., 2016), MapMan nomenclature developed for plant-specific pathways and processes (Navarro et al., 2019), and the Kyoto Encyclopedia of Genes and Genomes (KEGG) database for identification of overrepresented pathways (Kanehisa et al., 2016). Gene Ontology (GO) classification by biological processes of the identified proteins indicated that most of the proteins are involved in “Cellular (GO:0009987) and Metabolic (GO:0008152) processes.” This classification might be biased by the fact that these processes include the most abundant proteins, which are preferentially detected by MS, as observed in other proteomic analyses (Aroca et al., 2017; Huang et al., 2019) (Supplemental Fig. S8). An important subset of proteins is categorized within the GO “Response to stimulus” (GO:0050896), which includes ten secondary GO identifications, highlighting “Response to stress” and “Response to chemical” (Supplemental Fig. S8). GO classification of the proteins present only in wild type or *cas-c1* root hair samples, again showed that “Cellular and Metabolic processes” are the prevailing sets but within the GO classification “Biological regulation” (GO:0065007) there were 66 and 136 proteins in wild type and *cas-c1*, respectively (Supplemental Fig. S9). Among the proteins in this GO subclassification (Supplemental Table S1), we identified the transcription factor ELONGATED HYPOCOTYL 5 (HY5), a positive regulator of photomorphogenesis whose absence results in increased root hair length (Zhang et al., 2019), TRANSPARENT TESTA GLABRA 1 (TTG1), involved in trichome and root hair development (Tan et al., 2021), and ENHANCED ETHYLENE RESPONSE PROTEIN 5 (EER5), which functions in resetting the ethylene-signaling pathway (Christians et al., 2008). In this subset of proteins, we also identified in *cas-c1* but not in wild type, several proteins involved in hormone signal transduction such as the brassinosteroid co-receptor BRI1-ASSOCIATED RECEPTOR KINASE 1 (BAK1) and the BRASSINOSTEROID-SIGNALING KINASE 7 (BSK7), and auxin signaling and trafficking such as the AUXIN SIGNALING F-BOX2 (AFB2) and SHORT AND SWOLLEN ROOT 1 (NDP1/SSR1), which encodes a tetraticopeptide-containing repeat localized in mitochondria involved in root development (Han et al., 2021).

The identified proteins in both sample types were also analyzed based on their assigned functions and classified into 35 functional groups using the MapMan nomenclature (Navarro et al., 2019) (Fig. 6). The most abundant set corresponded to the general PROTEIN group, which included almost 20% of the total identified proteins in both cell type backgrounds with 853 and 898 elements in wild type and *cas-c1*, respectively. This bin set comprises several subsets involved in protein activation, protein synthesis and degradation, protein targeting, post-translational modification, protein folding and glycosylation (Supplemental Table S2). Taken into account the defect in development of the root hairs in *cas-c1*, we analyzed in detail the proteins involved in developmental processes in the protein subset present only in the samples of wild type and *cas-c1*. As shown in Supplemental Table S3, 7 and 16 proteins classified in development are only present in wild type and *cas-c1*, respectively. Among the proteins that are absent in *cas-c1* root hair-like cells, we found the protein kinase TARGET OF RAPAMYCIN (TOR), which functions as a central regulator of growth, coordinating nutritional and hormonal signaling, as well as a repressor of the autophagy process (Xiong and Sheen, 2014), EMBRYO DEFECTIVE 30 (EMB30), a GDP/GTP exchange factor for small G-proteins involved in the specification of apical-basal pattern formation and essential for cell division and expansion (Li et al., 2016a), and HSP4/SABRE protein, which mediates microtubule organization and affects root hair patterning (Pietra et al., 2015). Among the proteins that are only present in *cas-c1*, we identified PESCADILLO, which plays a role in root elongation and differentiation (Zografidis et al., 2014), and TRANSPARENT TESTA GLABRA 1 (TTG1), also identified in the GO classification of Biological regulation.

**Fig. 6.**
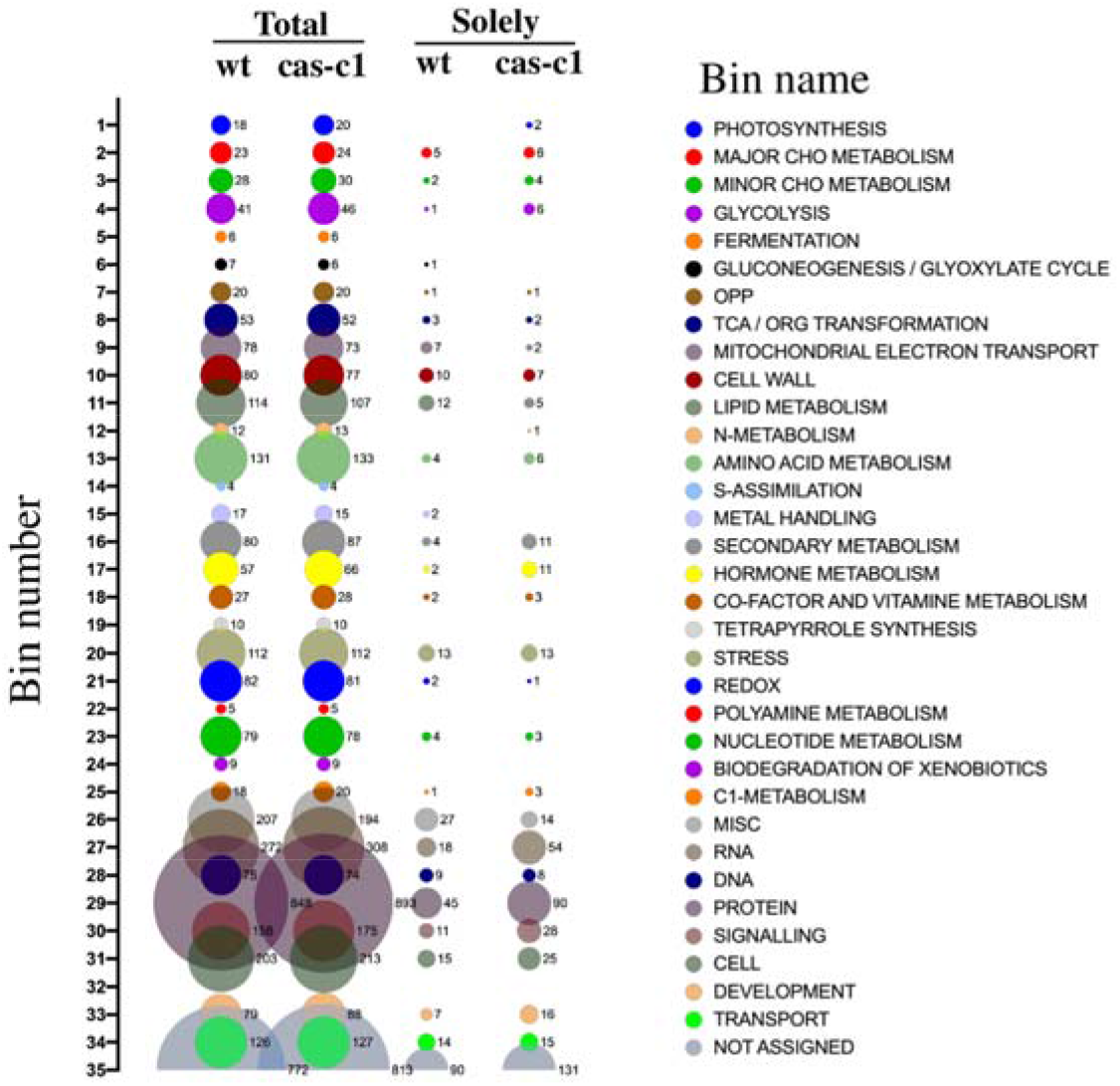
Bubble plot of the functional classification of the identified proteins according to the plant-specific database MapMan. Bubble columns represent the proteins classified within each bin of the total proteins identified in wild type (wt), *cas-c1* root hair samples, and those only present in wt or *cas-c1* root hair samples. Numbers besides the bubble represent the amount of proteins classified within the bin.

Finally, the proteins that were present only in wild type or *cas-c1* samples were also analyzed using the KEGG Search Pathway (Supplemental Table S4) to identify overrepresented pathways or processes. Apart from the Metabolic and Secondary metabolite biosynthetic pathways, the Spliceosome pathway is the one that shows the highest representation of proteins in *cas-c1* root hair samples with 22 proteins (Supplemental Table S5 and Supplemental Fig. S10). These 22 proteins include components of the small nuclear ribonucleoproteins U1, U2, U4, U5, U6 complex like SPF3 (At2g02570) and YLS8 (At5g08290); the Prp19 complex, and the Prp2 and Prp22 pre-mRNA-splicing factors RNA helicase. This is not surprising because the spliceosome is modified in root hair cells as compared to non-hair cells, and there is less intron retention in RH than in NH cells (Lan et al., 2013). In addition, the RNA transport and mRNA surveillance pathway are also overrepresented with 12 and 9 proteins respectively.

### *S*-cyanylation of root hair proteins

HCN can produce post-translational modification of proteins by *S*-cyanylation of cysteine residues (Garcia et al., 2019). *S*-cyanylation in peptides can be identified by a 25.0095 Da mass increase in the fragmentation spectrum. We identified 25 proteins *S*-cyanylated in wild type (Table 1) and 32 in *cas-c1* plants (Table 2), 8 of which are common in both. The low number of modified proteins identified is biased by the fact that a chemical selective method to label, enrich and detect *S*-cyanylated proteins is not available even though such does exist for other post-translational modifications including phosphorylation (Dudley and Bond, 2014), nitrosylation (Qu et al., 2016) or persulfidation (Aroca et al, 2017). Among the *S*-cyanylated proteins identified in *cas-c1* root hair samples, we observed four proteins of the methionine regeneration SAM cycle including COBALAMIN-INDEPENDENT METHIONINE SYNTHASE (MS1), METHIONINE SYNTHASE 2 (MS2), the S-ADENOSYL-L-HOMOCYSTEINE (SAH) HYDROLASE 2 (SAHH2) and the S-ADENOSYL-L-HOMOCYSTEIN HYDROLASE 1 (SAHH1/HOG1). These proteins have been previously detected as *S*-cyanylated in *cas-c1*, which indicates that the SAM cycle is a significant target process of HCN (Garcia et al., 2019). Fig. 7 shows the digestion pattern of the peptide containing the *S*-cyanylated Cys in SAHH1/HOG1. As shown in the table, a predicted ion containing Cys-CN is identified in the spectrum. Interestingly, *hog1* mutants show a root hair defective phenotype, similar to that of *cas-c1* (Wu et al., 2009).

**Table 1.**
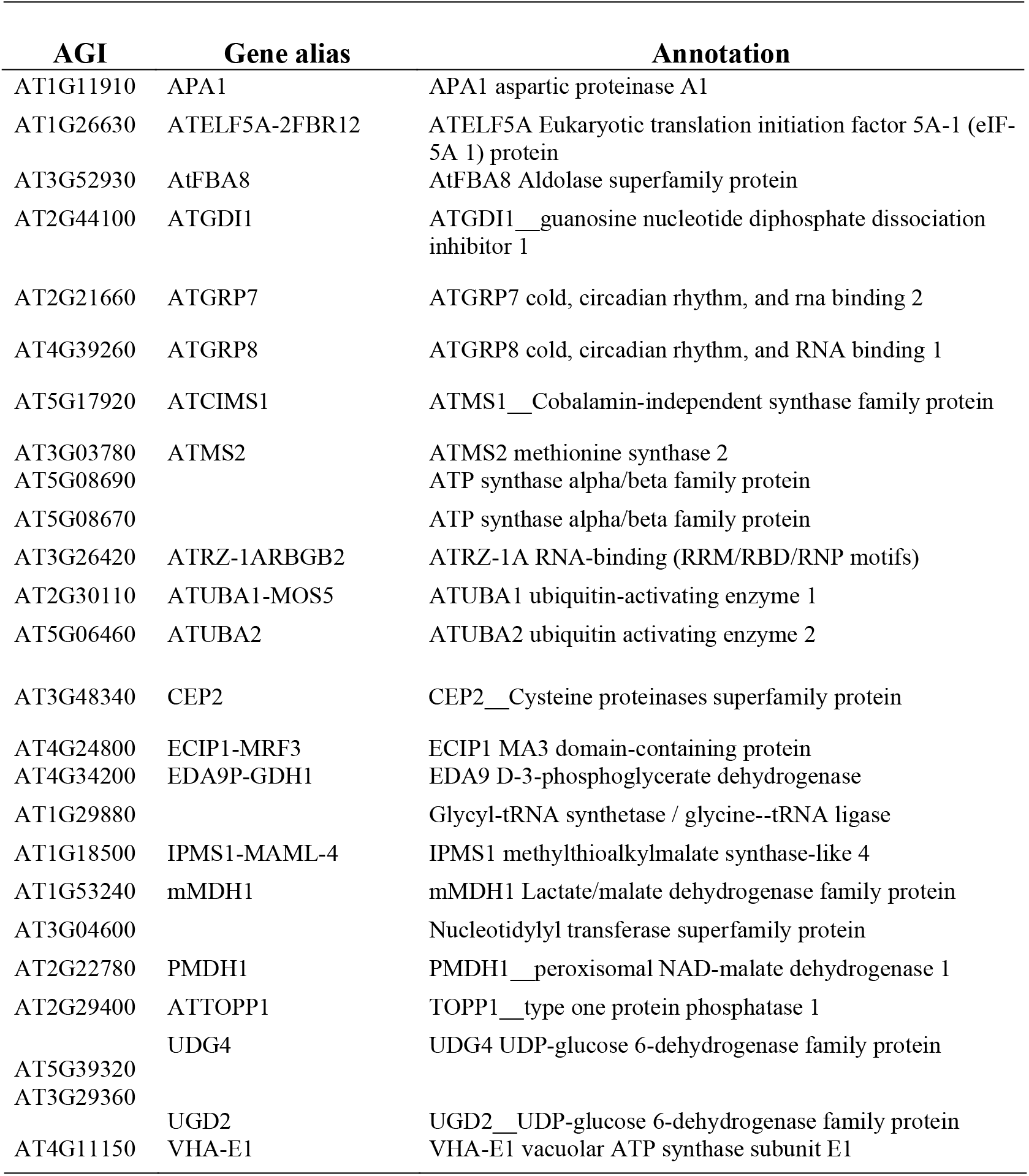
Cyanilated proteins identified in wild type root hair samples

| AGI | Gene alias | Annotation |
| --- | --- | --- |
| AT1G11910 | APA1 | APA1 aspartic proteinase A1 |
| AT1G26630 | ATELF5A-2FBR12 | ATELF5A Eukaryotic translation initiation factor 5A-1 (eIF-5A 1) protein |
| AT3G52930 | AtFBA8 | AtFBA8 Aldolase superfamily protein |
| AT2G44100 | ATGDI1 | ATGDI1__guanosine nucleotide diphosphate dissociation inhibitor 1 |
| AT2G21660 | ATGRP7 | ATGRP7 cold, circadian rhythm, and rna binding 2 |
| AT4G39260 | ATGRP8 | ATGRP8 cold, circadian rhythm, and RNA binding 1 |
| AT5G17920 | ATCIMS1 | ATMS1__Cobalamin-independent synthase family protein |
| AT3G03780 | ATMS2 | ATMS2 methionine synthase 2 |
| AT5G08690 |  | ATP synthase alpha/beta family protein |
| AT5G08670 |  | ATP synthase alpha/beta family protein |
| AT3G26420 | ATRZ-1ARBGB2 | ATRZ-1A RNA-binding (RRM/RBD/RNP motifs) |
| AT2G30110 | ATUBA1-MOS5 | ATUBA1 ubiquitin-activating enzyme 1 |
| AT5G06460 | ATUBA2 | ATUBA2 ubiquitin activating enzyme 2 |
| AT3G48340 | CEP2 | CEP2__Cysteine proteinases superfamily protein |
| AT4G24800 | ECIP1-MRF3 | ECIP1 MA3 domain-containing protein |
| AT4G34200 | EDA9P-GDH1 | EDA9 D-3-phosphoglycerate dehydrogenase |
| AT1G29880 |  | Glycyl-tRNA synthetase / glycine--tRNA ligase |
| AT1G18500 | IPMS1-MAML-4 | IPMS1 methylthioalkylmalate synthase-like 4 |
| AT1G53240 | mMDH1 | mMDH1 Lactate/malate dehydrogenase family protein |
| AT3G04600 |  | Nucleotidyl transferase superfamily protein |
| AT2G22780 | PMDH1 | PMDH1__peroxisomal NAD-malate dehydrogenase 1 |
| AT2G29400 | ATTOPP1 | TOPP1__type one protein phosphatase 1 |
| AT5G39320 | UDG4 | UDG4 UDP-glucose 6-dehydrogenase family protein |
| AT3G29360 | UGD2 | UGD2__UDP-glucose 6-dehydrogenase family protein |
| AT4G11150 | VHA-E1 | VHA-E1 vacuolar ATP synthase subunit E1 |

**Table 2.**
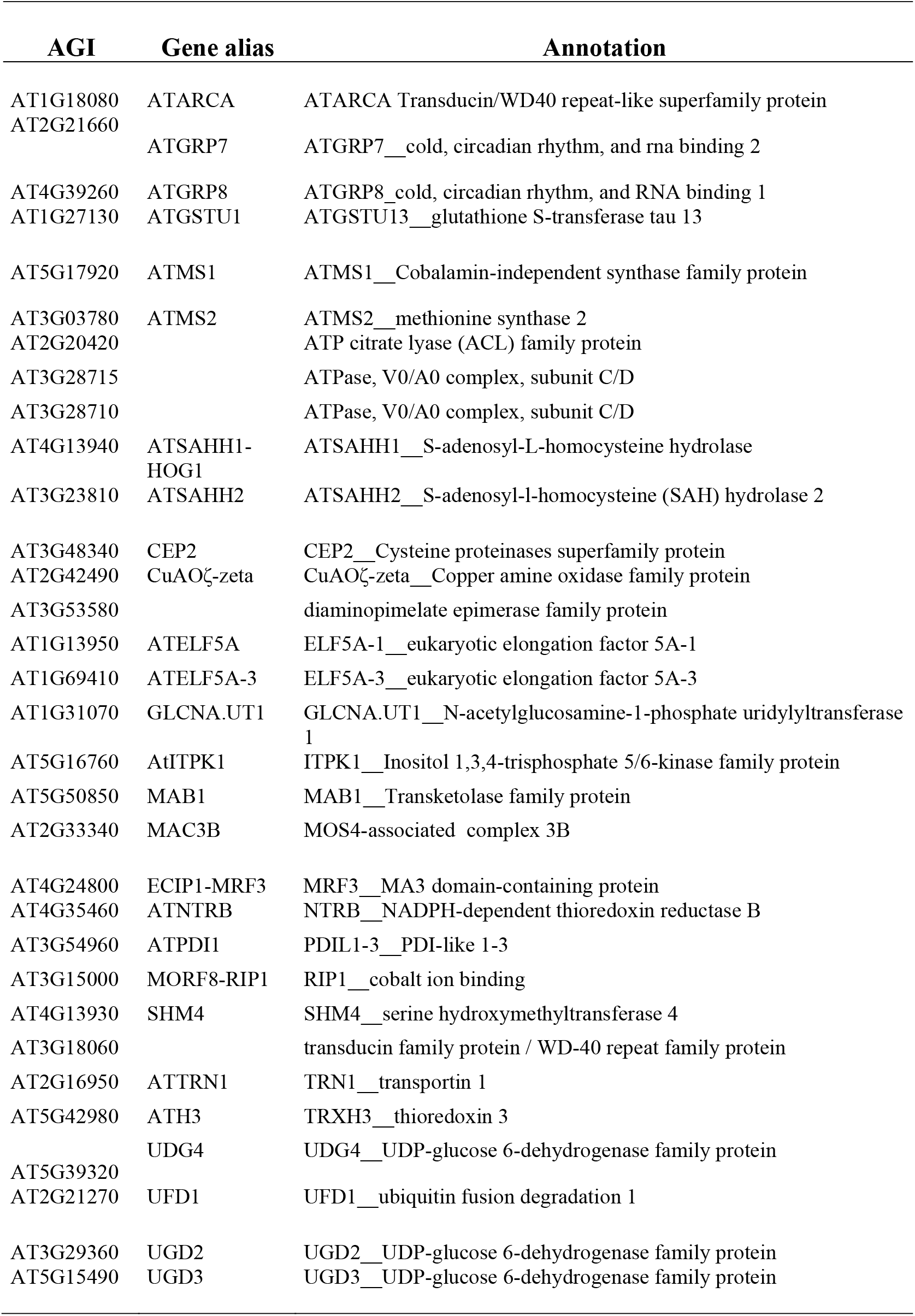
Cyanilated proteins identified in cas-c1 root hair samples.

**Fig. 7.**
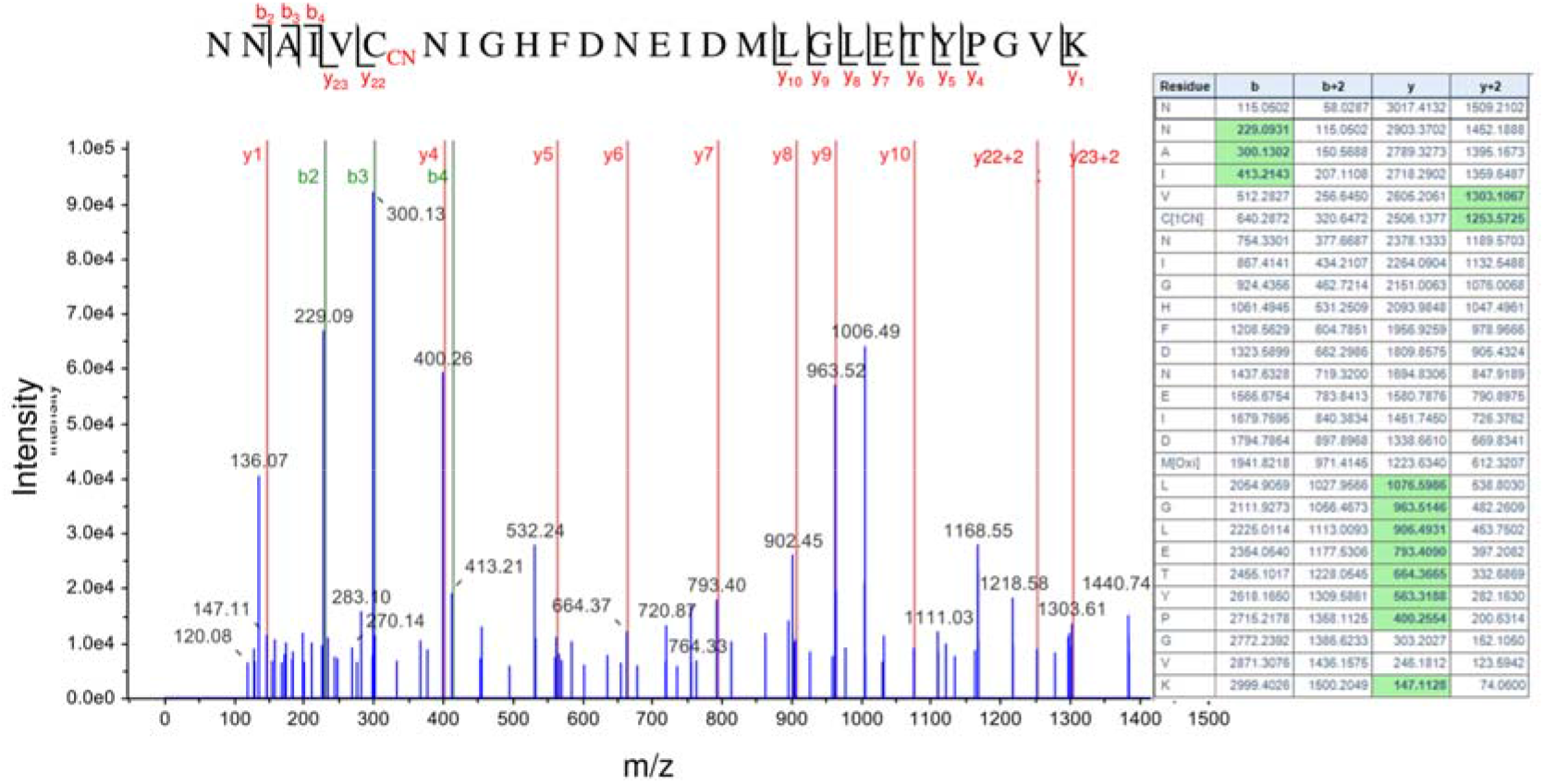
LC-MS/MS mass spectra of the tryptic peptide containing the *S*-cyanylated Cys^346^ residue of the S-Adenosyl homocysteine hydrolase SAHH1/HOG1 protein. The table contains the predicted ion type for the modified peptide, and the ions detected in the spectrum are highlighted in green color.

We also found that the GLYCINE-RICH RNA BINDING PROTEIN 7 (GRP7) is *S*-cyanylated in the two sets of proteins. Fig. 8 shows the digestion pattern of the peptide containing the *S*-cyanylated Cys in GRP7 and predicted ions containing Cys-CN are identified in the spectrum. GRP7 binds FER and is phosphorylated in the RAPID ALKALINIZATION FACTOR PEPTIDES (RALF)-FER-GRP7 complex, which is implicated in RNA alternative splicing during root hair formation (Wang et al.). This, together with the *in silico* data obtained in a previous section, supports a relationship between *cas-c1*/HCN and *feronia*.

**Fig. 8.**
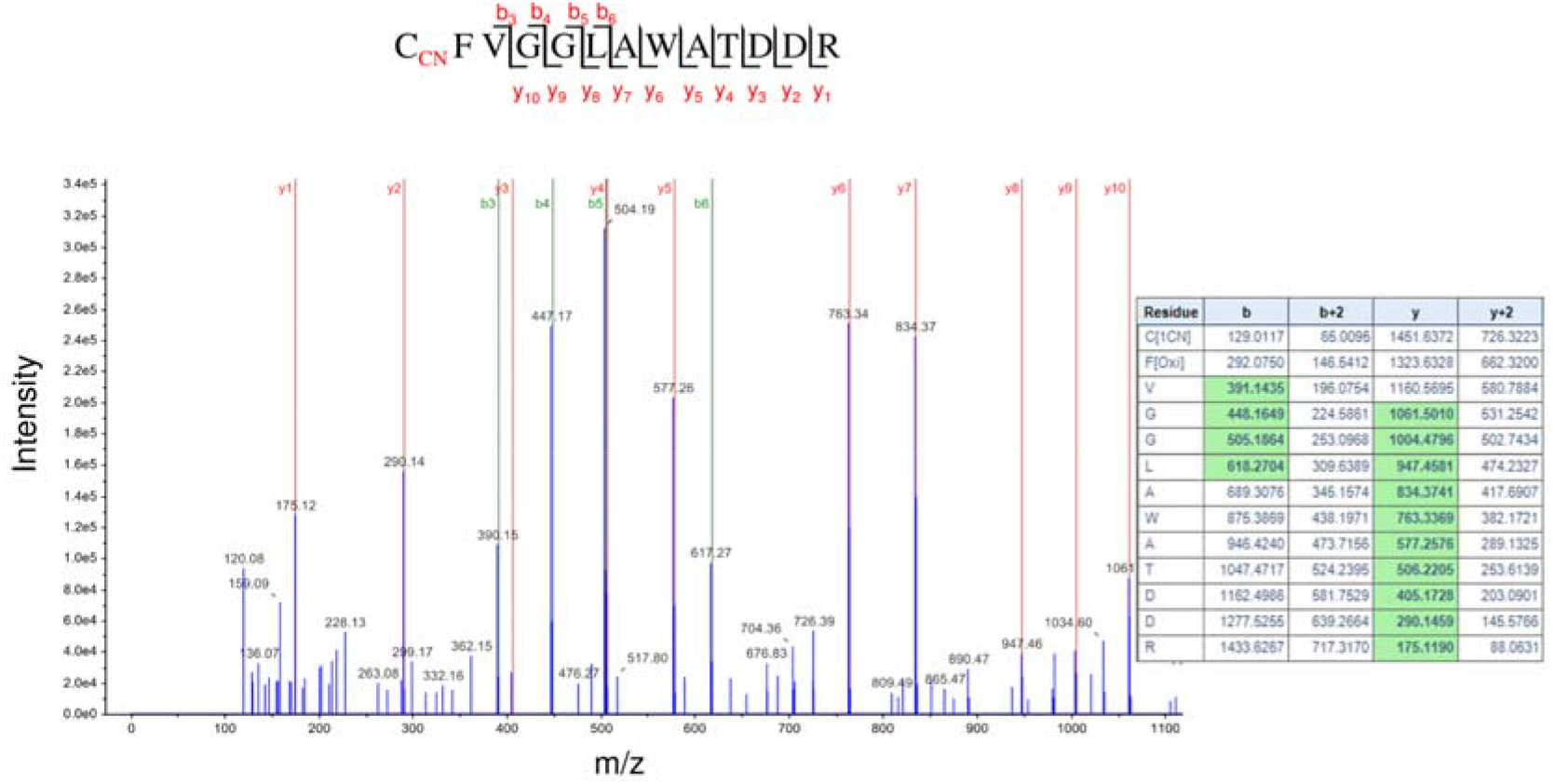
LC-MS/MS mass spectra of the tryptic peptide containing the *S*-cyanylated Cys^10^ residue of the Glycine Rich Protein 7, (GRP7). The table contains the predicted ion type for the modified peptide, and the ions detected in the spectrum are highlighted in green color.

## Discussion

The role of ROS, particularly O_2_^•−^ and H_2_O_2_, in root and root hair development has been extensively studied. These species exhibit different gradient distributions in the roots (Yamada et al.; Zhou et al., 2020). A gradient of O_2_^•−^ is essential for root hair elongation and this gradient at the root hair tip is lacking in the *cas-c1* mutant (Garcia et al., 2010). On the other hand, a transcriptomic analysis suggested that *CAS-C1* might play a role in QC, phloem or developing cortex formation or function (Lan et al., 2013). Despite the fact that *cas-c1* plants have a higher ROS content than wild type in control conditions (Garcia et al., 2013), no visible differences were detected in the QC, developing and maturation zone morphology or root length in *cas-c1* as compared to wild type. In the root maturation zone, the epidermal cells located in the H cell position emerge to form hair cells, and plant hormones, such as auxin and ethylene regulate root hair formation *via* ROS. Moreover, superoxide anion drives the elongation of the root hair *via* NADPH oxidase action (Carol and Dolan, 2006). However, the *cas-c1* phenotype is not related to ROS even though there is a possible direct inhibition of the NADPH oxidase RHD2/ATRBOHD by HCN (Arenas-Alfonseca et al., 2018a, 2018b). Thus, the proteomic analysis performed on isolated root hair cells that is described here provides additional functional data for the CAS-C1 protein and HCN role in root hair development.

Previously, there has been little proteomic analysis performed on root hair cells (Petricka et al., 2012) (Lan et al., 2013). In this work, more than 3700 proteins were identified in each sample, of which 457 were identified in *cas-c1* but not in wild type. This demonstrated that the suppression of CAS-C1 activity and the consequent accumulation of HCN in the *cas-c1* mutant generates important changes in the root hair proteome (Garcia et al., 2010; Arenas-Alfonseca et al., 2018a, 2018b). The analysis of these 457 proteins using the KEGG database showed an over-representation of proteins involved in the spliceosome, the RNA transport and mRNA monitoring pathways. The spliceosome is a large complex formed by small nuclear ribonucleoproteins (snRNPs) (Lorković et al., 2000) that process introns in pre-mRNA to convert it into mature mRNA, a fundamental stage in gene expression in eukaryotes. RNA binding proteins, RNA dependent ATPases, RNA helicases and protein kinases are also involved in this process. Many of the spliceosome proteins, identified exclusively in *cas-c1*, participate in various developmental processes, including flowering, apical dominance, leaf and rosette size, shape and morphology (Lopato et al., 1999; Ru et al., 2008). An in-depth study of gene expression in specific root cells showed great variation in the presence of RNAs with non-coding intergenic regions that contribute to the functional specialization of the cell, demonstrating that alternative intron processing serves to regulate differentiation (Li et al., 2016b). Therefore, RH display less intron retention than NH and this proteomic difference between *cas-c1* and wild type plants is consistent with this finding.

It is noteworthy that we only detected TRANSPARENT TESTA GLABRA1 (TTG1) in *cas-c1* root hairs. This protein, together with WEREWOLF (WER) and GLABRA3 (Gl3), form a complex in non-hair cells that induces GLABRA2 (GL2) expression and controls non-root hair cell fate by the GL2-mediated suppression of a set of root hair specific transcription factors (Shibata and Sugimoto, 2019). Therefore, a *ttg1* knockout mutant displays a hairy phenotype. The presence of abnormally high amounts of TTG1 in the *cas-c1* root hairs could result in a hairless phenotype. However, we demonstrate that the *cas-c1* mutation does not affect root hair cell fate as a COBL9-GFP fusion locates exclusively in hair cells and follows the wild-type alternate H-NH pattern, and COBL9 action locates downstream of TTG1 in the genetic pathway that leads to root hair specification and elongation (Grierson and Schiefelbein, 2009).

Previous genetic experiments demonstrated that CAS-C1 acts between SUPERCETIPEDE 1 (SCN1), a Rho GTPase GDP dissociator inhibitor, which acts as a negative regulator of ROPs, and the NADPH oxidase action, which is regulated by ROPs (Arenas-Alfonseca et al., 2018b, 2018a). This points to ROPs as possible targets of cyanide action. FERONIA regulates ROP-signaled root hair development (Duan et al., 2010), and the transcriptional profile of its mutant shows a significant number of mis-regulated genes common to mis-regulated proteins in *cas-c1*. Thus, we can hypothesize that both *fer* and *cas-c1* share common elements in the root hair development process or that one acts upstream of the other. This may involve *S*-cyanylation of proteins regulated by FER in RH elongation as a mode of action of HCN. Candidates could be peroxidases, ROPs or other proteins involved in RH elongation mediated by FER. This opens an interesting focus on the interactions between *cas-c1*, *feronia* and *S*-cyanylation that will be the subject of future research. On the other hand, it has been demonstrated that the interaction between the peptide rapid alkalinization factor (RALF) and FERONIA modulates protein synthesis by the phosphorylation of a translation initiation factor (Zhu et al., 2020). Several initiation factors have been identified in samples of both wild-type and *cas-c1* root hairs. The translation initiation factor 2 is present only in *cas-c1* root hairs, as well as another putative initiation factor, thus establishing a possible relationship between FERONIA and CAS-1 at the functional level. This is consistent with the MapMan analysis, in which approximately 20% of the genes belong to PROTEIN group, opening also the question of whether proteins are affected at the post-translational level by HCN and its associated post-translational modification, the *S*-cyanylation (Garcia et al., 2019). Moreover, GRP7 is *S*-cyanylated in root hairs, being a protein that works together with the RALF-FER module to adjust the alternative RNA splicing in response to environmental and developmental factors (Wang et al.). Therefore, HCN signaling and root hair elongation could be coupled by the *S*-cyanylation of this RNA-binding protein and its interaction with FERONIA.

Among the proteins that are *S*-cyanylated in *cas-c1* plants, we found a significant number of members of the SAM cycle, a relevant finding because in our previous work we also found these proteins in *cas-c1* samples (Garcia et al., 2019). The SAM cycle is involved in DNA methylation through SAM, a methylation DNA agent. DNA methylation is a conserved epigenetic marker that regulates many developmental and stress responses and adaptation in plants, including the transmission of a stress memory that can be important in the response to pathogens. Mutant lines (sahh1/hog1) with reduced levels of the S-ADENOSYL-L-HOMOCYSTEIN HYDROLASE 1 enzyme show a deficiency in root hair development pointing to a relationship between methylation level and RH development (Wu et al., 2009). There is substantial evidence that correlates the levels of SAM cycle components with DNA methylation changes (Zhang, 2018; Gonzalez and Vera, 2019). In plants, inhibition of the METHIONINE SYNTHASE interferes with the immune response of the plant to pathogens. It has been demonstrated extensively that *cas-c1* plants are more resistant to biotrophic pathogens (Garcia et al., 2013; Garcia et al., 2014; Arenas-Alfonseca et al., 2021). In fact, very low concentrations of KCN (1 *μ*M) are able to induce the plant response to pathogens and induce the *PATHOGENESIS-RELATED 1* (*PR1*) gene (Arenas-Alfonseca et al., 2021), and it has been very recently described that this fact is not exclusive to plants. In mammalian cells, very low concentrations of HCN (from nM to 1 *μ*M) induce cellular proliferation and bioenergetics *via* cytochrome C oxidase stimulation (Randi et al., 2021). Understanding the participation of HCN in regulating the SAM cycle and DNA methylation in plant priming is one of our future goals.

## Materials and methods

### Plant material and growth conditions

Arabidopsis (*Arabidopsis thaliana*) wild type ecotype Col-0 and the *cas-c1* T-DNA insertion mutant (*cas-c1*; SALK_103855; (Garcia et al., 2010)) were grown in soil or vertical positioned plates in a photoperiod of 16 h of white light (120 *μ*mol m^−2^ s^−1^) at 20°C and 8 h of dark at 18°C. Seeds were surface sterilized using 50 % (vol/vol) bleach and 0.1% Tween 20 (Sigma) for 15 min and then rinsed five times with sterile water. All seeds were plated on standard MS media (1× Murashige and Skoog salt mixture, Caisson Laboratories), 0.5 g/L MES, 1% Sucrose, and 1% Agar (Difco) and adjusted to pH 5.7 with KOH. All plated seeds were stratified at 4 °C for 2 d before germination.

For nutrient deficiency assays, 1x Murashige and Skoog lacking sulphur, nitrogen or phosphate (Caisson Laboratories) were used.

### Root meristem visualization

For root meristem analysis, wild type and *cas-c1* mutant plants were grown for 7 days in vertical MS medium plates. Roots were then mounted in 10 mg/mL propidium iodide (PI) in water and observed using a 20 X objective with a Zeiss LSM 880 laser scanning confocal microscope. Excitation and detection window were set as follows: excitation at 561 nm and detection at 570-650 nm.

### Expression of pCOBL9:GFP in roots

The pCOBL9:GFP construct (Brady et al., 2007a; Petricka et al., 2012) was introduced into wild type (Col) and *cas-c1* mutant plants. Two independent T3 lines in each background were grown for 6 days in MS media. Roots from transgenic plants were imaged using a Leica TCS SP2 spectral confocal microscope. Samples were excited using an argon ion laser at 488 nm; emission was detected between 510 and 580 nm for GFP imaging (pseudocolored green). The microscopy images were processed using Leica Confocal Software.

### Root protoplasts isolation

500-1000 wild type and *cas-c1* mutant seeds were grown for 6 days on Nitex nylon mesh MS vertical plates (100mm x 100mm x15mm). Roots were sliced and placed in a petri dish (Falcon 351008) holding one 70 *μ*m strainer (Falcon 352350) with 7 ml buffer B (cellulase 15 g l^−1^, (Sigma #C1794) and pectolyase 1 g l^−1^, (Sigma #P3026) in buffer A). Roots were incubated for 1 h at room temperature and 85 rpm. The filtered liquid was transfered to 15 ml Falcon and centrifuged for 6 min at 22 °C, 200 x g. Supernatant was then removed and the pellet was resuspended in 300 *μ*l of buffer A (mannitol 600 mM, MgCl_2_ 2 mM, CaCl_2_ 2 mM, MES 2 mM, KCl 10 mM and BSA 0,1% (p/v), pH 5.5). The mix was after filtered in a 70 and 40 *μ*m (Falcon 352340) strainer, respectively, and the filtered cells were kept in a 5 ml polystyrene tube.

### Fluorescent activated cell sorting (FACS)

Protoplasts from wild type pCOBL9:GFP and *cas-c1* pCOBL9:GFP transgenic plants were selected in a sorter MoFlo Astrios EQ (Beckman-Coulter), with a 70 *μ*m nozzle at a rate of 5,000 to 10,000 events per second and a fluid pressure of 60 psi. Protoplast of non-transgenic wild type and *cas-c1* plants were used as negative control. GFP positive cells were based on the negative control. They were read using a bandpass filter of 526/52. Protoplast were then collected in PBS buffer (NaCl 80 g l^−1^, KCl 2 g l^−1^, Na_2_HPO_4_ 14.4 g l^−1^ y KH_2_PO_4_ 2.4 g l^−1^, pH 7.4), frozen in liquid nitrogen and stored for protein extraction.

### LC-MS/MS based root hair proteomics

Protein extraction from sorted protoplast was done and protein amount was measured by Bradford assay (Bradford, 1976) and normalized with 50 mM NH_4_HCO_3_ pH 8.0 to 0.1-1 *μ* g *μ* l^−1^. RapiGest SF Surfactant (Waters) 0.2% (v/v) was added. Proteins were incubated 10 min at 40 °C, 100 rpm and reduced with DTT 10 mM. After, proteins were incubated again at 10 °C for 40 min and 100 rpm. Samples were cool down at room temperature and alkylated with iodoacetamide for 30 min at RT. Protein digestion was done with trypsin (Promega) at a 1:50 ratio and incubated for 4 h. Peptides were dried in a vacuum centrifuge. Prior to LC-MS/MS analysis, samples were resusupended in TFA 1 % (v/v), acetonitrile 2 % (v/v). Samples were then incubated for 2h at 60 °C and 100 rpm and centrifugated at 15000 rpm for 5 min. Supernatant was stored for LC-MS/MS. 1 *μ* g of protein was analyzed in a LC-MS/MS elution gradient for 90 min in a mass spectrometer “Q Exactive HF-X Hybrid Quadrupole-Orbitrap”, (Thermo Scientific). LCMS and MS/MS data was process by Proteome Discoverer 2.2 program (Thermo Scientific) and analyzed for identification in MASCOT (Matrix Science) against TAIR10. Search tolerances were 5 ppm for precursor ions and 0.02 Da for product ions using trypsin specificity with up to two missed cleavages. All searched spectra were imported into Scaffold (v4.3, Proteome Software) and scoring thresholds were set to achieve a peptide false discovery rate of 1% using the PeptideProphet algorithm.

### Proteomic analysis

Protein function analysis and classification was performed using three biological gene/protein databases: MapMan (Klie and Nikoloski, 2012), Uniprot (Bairoch et al., 2005; Pundir et al., 2016) and the Kyoto Encyclopedia of Genes and Genomes (KEGG) (Kanehisa et al., 2016).

### Accession Numbers

Sequence data from this article can be found in the website http://www.arabidopsis.org/ for Arabidopsis genes with locus identifiers provided in this study. The mass spectrometry proteomics data have been deposited in the ProteomeXchange Consortium *via* the PRIDE partner repository with the data set identifier PXD028991 (Vizcaíno et al.).

## Acknowledgements

We acknowledge Dr. Philip N. Benfey and Duke University for accepting L.A.-A. for a short-stay to perform the FACS and proteomics experiments, and Dr. Philip N. Benfey for his valuable suggestions in both work development and manuscript writing. We are grateful to Dr. Heather Belcher for her help with root protoplast protocol. We acknowledge the European Regional Development Fund, Ministerio de Economía y Competitividad, the Agencia Estatal de Investigación and CSIC (BIO2016-76633-P and 201840I085 grants) and the Junta de Andalucía (P20_00030 grant) for funding. L.A.-A. thanks the Ministerio de Economía y Competitividad for fellowship support through the program of Formación de Personal Investigador.

The authors have no conflict of interest to declare.

